# *Zobellia alginoliquefaciens* sp. nov., a new flavobacteria isolated from the epibiota of the brown alga *Ericaria zosteroides* (C.Agardh) Molinari & Guiry 2020

**DOI:** 10.1101/2023.03.13.532333

**Authors:** Tristan Barbeyron, Nolwenn Le Duff, Eric Duchaud, François Thomas

## Abstract

Strain LLG6346-3.1^T^, isolated from the thallus of the brown alga *Ericaria zosteroides* collected in Mediterranean Sea near Bastia in Corsica, France, was characterized using a polyphasic method. Cells were Gram-stain-negative, strictly aerobic, non-flagellated, motile by gliding, rod-shaped and grew optimally at 30-33 °C, at pH 8-8.5 and with 4-5 % NaCl. Strain LLG6346-3.1^T^ used the seaweed polysaccharide alginic acid as sole carbon source which was vigorously liquefied. Phylogenetic analyses showed that the bacterium is affiliated to the genus *Zobellia* (family *Flavobacteriaceae*, class *Flavobacteriia*). Strain LLG6346-3.1^T^ exhibited 16S rRNA gene sequence similarity values of 98.5 and 98.3 % to the type strains of *Zobellia russellii* and *Zobellia roscoffensis* respectively, and of 97.4-98.2 % to other species of the genus *Zobellia*. The DNA G+C content of strain LLG6346-3.1^T^ was determined to be 38.28 mol%. Digital DNA-DNA hybridization predictions by the ANI and GGDC methods between strain LLG6346-3.1^T^ and other members of the genus *Zobellia* showed values of 76-88 %, and below 37 %, respectively. The phenotypic, phylogenetic and genomic analyses show that strain LLG6346-3.1^T^ is distinct from species of the genus *Zobellia* with validly published names and that it represents a novel species of the genus *Zobellia*, for which the name *Zobellia alginoliquefaciens* sp. nov. is proposed. The type strain is LLG6346-3.1^T^ (RCC 7657^T^ = LLG 32918^T^).

---

The genus *Zobellia* belongs to the family *Flavobacteriaceae* (order *Flavobacteriales*, class *Flavobacteriia*), was proposed by Barbeyron et al. [1] with *Zobellia galactanivorans* as the type species of the genus. At the time of writing, the genus *Zobellia* comprises 8 validly named species, all isolated from marine environments and mostly from macroalgae. For example, *Z. galactanivorans* Dsij^T^ was retrieved as an epibiont of the red alga *Delesseria sanguinea* [2], *Z. russellii* KMM 3677^T^ and *Z. barbeyronii* 36-CHABK-3-33^T^ were isolated from the green algae *Acrosiphonia sonderi* and *Ulva* sp. respectively [3, 4], while *Z. laminariae* KMM 3676^T^ originated from the brown alga *Saccharina japonica* [3] and *Z. nedashkovskayae* Asnod2-B07-B^T^ and *Z. roscoffensis* Asnod1-F08^T^ from the brown alga *Ascophyllum nodosum* [5]. Moreover, metagenomics survey show that the genus *Zobellia* is part of the microbiota of healthy macroalgae [6, 7]. Recent development of *Zobellia*-specific quantitative PCR primers and FISH probes confirmed the presence of the genus on the surface of diverse macroalgal species, with ca. 10^3^-10^4^ 16S rRNA copies.cm^-2^ [8]. Strain LLG6346-3.1^T^ was isolated in May 2019 from the surface of *Ericaria zosteroides* (C.Agardh) Molinari & Guiry 2020 thallus during a sampling campaign in the Mediterranean Sea near Negru in Corsica (France, GPS 42.769040 N, 9.333530 E). The algal specimen was collected manually by divers at ca. 20 m depth, before swabbing in the laboratory and inoculation on ZoBell 2216E-agar plates [9]. Here, we present a detailed taxonomic investigation of strain LLG6346-3.1^T^ using a polyphasic approach, including some genomic data deduced from its complete genome and also techniques of whole-genome comparison such as the Average Nucleotide Identity (ANI) and dDDH (digital DNA–DNA hybridization).

For comparison, *Zobellia russellii* KMM 3677^T^ = LMG 22071^T^ [3] purchased from the Collection de l’Institut Pasteur (CIP; France) and *Zobellia roscoffensis* Asnod1-F08^T^ = CIP 111902^T^ = RCC 6906^T^ [5] isolated in our laboratory, were used as related type strains., *Z. russellii* KMM 3677^T^ and *Z. roscoffensis* Asnod1-F08^T^ were studied in parallel with strain LLG6346-3.1^T^ for all phenotypic tests except for the temperature, pH and NaCl ranges of growth. The three strains were routinely cultivated on ZoBell medium 2216E, either liquid or solidified with 1.5 % (w/v) agar. Pure cultures were stored at −80 °C in the culture medium containing 20 % (v/v) glycerol. All experiments were performed in triplicate. Assays of optimal temperature, pH and NaCl concentration were performed in 24-well plate containing 600 μl of medium inoculated with 12 μl of an overnight preculture. Optical density at 600 nm was measured in a Spark Tecan plate reader. The plate lid was pre-treated with 0.05 % Triton X-100 in 20 % ethanol to avoid condensation [10]. Growth was evaluated in ZoBell broth at 4, 13, 20, 24, 27, 30, 33, 36, 37, 38 and 40 °C. The optimal pH value for growth was determined at 30 °C in ZoBell broth with pH values adjusted by using 100 mM of the following buffers: MES for pH 5.5; Bis Tris for pH 6.0, 6.5, 7.0 and 7.5; Tris for pH 8.0, 8.5 and 9.0; CHES for pH 9.5 and CAPS for pH 10.0, 10.5 and 11.0. The effect of NaCl on growth was determined at 30 °C and at pH 8 in ZoBell broth prepared with distilled water containing 0, 0.5, 1.0, 2.0, 3.0, 3.0, 4.0, 5.0, 6.0, 7.0, 8.0, 9.0 and 10.0 % NaCl.

Cell morphology and motility were investigated on wet mounts of an exponential phase ZoBell broth culture at 20 °C, by using phase-contrast microscopy on an Olympus BX60 instrument (Olympus, Tokyo, Japan). The Ryu non-staining KOH method [11] was used to test the Gram reaction.

Oxidase activity was assayed using discs impregnated with N,N,N’,N’-tetramethyl-p-phenylenediamine dihydrochloride reagent (bioMérieux). Catalase activity was assayed by mixing one colony from a ZoBell agar plate with a drop of hydrogen peroxide (3 %, v/v). Nitrate reductase activity was assayed using ZoBell broth containing 10 g l^-1^ of sodium nitrate. Nitrate reductase activity was revealed after growth at 20 °C and addition of Griess Reagent. Amylase activity was assayed on 0.2 % (w/v) soluble starch ZoBell agar plates. DNase activity was detected on DNA agar (Difco) prepared with seawater. Amylase and DNase activities were revealed by flooding the plates with Lugol’s solution or 1 M HCl, respectively. The degradation of Tween compounds (1 %, v/v) was assayed in ZoBell agar according to Smibert & Krieg [12]. Agarase, κ-carrageenase and ι-carrageenase activities were tested by inoculating ZoBell media solidified with (per litre): 15 g agar (Sigma-Aldrich, ref. A7002), 10 g κ-carrageenin (X-6913; Danisco) or 20 g ι-carrageenin (X-6905; Danisco) respectively. Alginate lyase activity was tested by inoculating ZoBell media solidified with 10 g l^-1^ sodium alginate (Sigma-Aldrich, ref. 180947) according the Draget’s method [13]. Strains were considered positive when colonies liquefied or produced craters in the solidified substrate. Additional phenotypic characterizations were performed using API 20 E, API 20 NE, API 50CH and API ZYM strips according to the manufacturer’s instructions (bioMérieux) except that API AUX medium and API 50 CHB/E medium were adjusted to 2.5 % NaCl. All strips were inoculated with cell suspensions in artificial seawater and incubated at 20 °C for 72 h. The ability to use carbohydrates as sole carbon and energy sources was also tested in marine minimal medium [14] containing 2.5 g l^-1^ of the following sugars (all from Sigma-Aldrich unless otherwise stated): D-glucose, D-galactose, D-fructose, L-rhamnose, L-fucose, D-xylose, L-arabinose, D-mannose, sucrose, lactose, maltose, D-mannitol, D-raffinose, amylopectin (Merck), arabinan from sugar beet (Megazyme), arabinoxylan from wheat (Megazyme), xylan from beechwood, pectin from apple, agar, porphyrin from *Porphyra* sp. (home-made extract) laminarin (Goëmar), galactan from gum arabic, galactomannan from carob (Megazyme), glucomannan from konjac (Megazyme), alginic acid from *Laminaria digitata* (Danisco), ι-carrageenin from *Euchema denticulatum* (Danisco), κ-carrageenin from *Euchema cottonii*,λ-carrageenin (Dupont), lichenin (Megazyme), ulvin from *Ulva* sp. (Elicityl), xyloglucan from tamarind seed (Megazyme) and sulphated fucoidin from *Ascophyllum nodosum* (kindly provided by Algues et Mer) and *Laminaria hyperborea* (home-made extract).

Sensitivity to antibiotics was tested by the disc-diffusion method on ZoBell agar plates and using antibiotic discs (Bio-Rad) containing (μg per disc, unless otherwise stated): penicillin G (10 IU), ampicillin (10), carbenicillin (100), oxacillin (5), streptomycin (500), kanamycin (30), chloramphenicol (30), tetracycline (30), lincomycin (15), bacitracin (130), rifampicin (30), vancomycin (30). The effects of the antibiotics on cell growth were assessed after 24 h of incubation at 20 °C, and susceptibility was scored on the basis of the diameter of the clear zone around the disc.

Genomic DNA was extracted from 500 μl of culture of strain LLG6346-3.1^T^ in Zobell 2216E broth using the Genomic-tip 20/G kit (Qiagen) following the manufacturer’s instructions. The Illumina sequencing library was prepared using the Nextera XT DNA kit (Illumina) and sequenced using Illumina MiSeq v3 PE300, resulting in 4,268,034 quality-filtered reads (Table S1). The Nanopore sequencing library was prepared using Ligation Sequencing Kit 1D (SQK-LSK109) and sequenced using MinION flow cell R9.4.1, resulting in 100,270 reads of average length 22,935 nt. Hybrid assembly was performed using unicycler v 0.4.8 in conservative mode and otherwise default settings [15]. The 16S rRNA gene sequence was amplified by PCR using pure genomic DNA as template and primer pairs specific for Bacteria, 8F [16] and 1492R [17]. PCR reactions were typically carried out in a volume of 20 μl containing 10-100 ng template, 0.2 μM each specific primer, 200 μM each dNTP, 1X GoTaq buffer (Promega) and 1.25 U GoTaq DNA polymerase (Promega). PCR conditions were as follows: initial denaturation for 10 min at 95°C, followed by 35 cycles of 1 min at 95°C, 30 sec at 50°C, 2 min at 72°C, and final extension of 5 min at 72°C. PCR products were purified using the ExoSAP-IT Express kit according to the manufacturer’s protocol (ThermoFisher Scientific) and sequenced by using BigDye Terminator V3.1 (Applied Biosystems) and an ABI PRISM 3130xl Genetic Analyzer automated sequencer (Applied Biosystems/Hitachi). Chargal?s coefficient of the genomic DNA of strain LLG6346-3.1^T^ was deduced from the complete genome sequence and expressed as the molar percentage of guanine + cytosine. The nucleotide sequence of the 16S rRNA gene deduced from the complete genome sequence of the strain LLG6346-3.1^T^ and sequences of the 16S rRNA genes from all valid species of the genera *Zobellia*, *Maribacter* and some other related genera of the *Flavobacteriaceae* family were aligned using the software MAFFT version 7 with the L-INS-I strategy [18]. The alignment was then manually refined and phylogenetic analyses, using the neighbour-joining [19], maximum-parsimony [20] and maximum-likelihood [21] methods, were performed using the MEGA 6 package [22]. The different phylogenetic trees were built from a multiple alignment of 50 sequences and 1437 positions. For the neighbour-joining algorithm, the evolutionary model Kimura Two Parameters [23] was used. The maximum-likelihood tree was calculated using the evolutionary model GTR (Generalised Time Reversible) [24] with a discrete Gamma distribution to model evolutionary rate differences among sites (4 categories). This substitution model was selected through submission of the alignment to the online server IQ-TREE (http://iqtree.cibiv.univie.ac.at/). The maximum-parsimony tree was obtained using the Subtree-Pruning-Regrafting algorithm [24]. Bootstrap analysis was performed to provide confidence estimates for the phylogenetic tree topologies [25]. A phylogenomic tree was performed on the web server M1CR0B1AL1Z3R [26]. Briefly, a total of 768 conserved orthologous ORFs were detected (identity > 80%, e-value < 0.001). Sequences were aligned using MAFFT [18] and maximum-likelihood [21] phylogeny was reconstructed using RaxML [27] with 100 bootstrap iterations. Pairwise comparisons of 16S rRNA gene sequences were made by using the database EzBioCloud (https://www.ezbiocloud.net/identify) [28]. Genomic relatedness was investigated by comparing the strain LLG6346-3.1^T^ genome sequence with those of the type strains of other *Zobellia* species using the Average Nucleotide Identity (ANI; http://jspecies.ribohost.com/jspeciesws/#analyse) [29–31] and the dDDH via the online server Genome to Genome Distance Calculator 2.1 (GGDC; http://ggdc.dsmz.de/distcalc2.php) [32]. The results from GGDC analysis were obtained from the alignment method Blast+ and the formula 2 (sum of all identities found in HSPs / by overall HSP length) for incomplete genome sequences [33, 34]. Exploration of carbohydrate active enzyme-coding genes in the genomes of the strains LLG6346-3.1^T^, *Z. russellii* KMM 3677^T^ and *Z. roscoffensis* Asnod1-F08^T^ was carried out via the online server Microscope from the French National Sequencing Center (http://www.genoscope.cns.fr/agc/microscope/mage) [35] and the CAZy database (www.cazy.org) [36].

The best pairwise comparison score with 16S rRNA gene from the strain LLG6346-3.1^T^ (1516 bp) were obtained with *Zobellia russellii* KMM 3677^T^ (98.6 %) (Table S1, available in the online version of this article). Phylogenetic analyses by neighbour-joining of 16S rRNA genes from species of the family *Flavobacteriaceae* showed that strain LLG6346-3.1^T^ belongs to the genus *Zobellia* and forms a clade with strains *Z. russellii* KMM 3677^T^ and *Z. roscoffensis* Asnod1-F08^T^ (Fig 1). The 16S rRNA gene sequence similarities between the strain LLG6346-3.1^T^ and other *Zobellia* species were in the range of 97.4 % with *Z. barbeyronii* 36-CHABK-3-33^T^ and *Z. nedashkovskayae* Asnod2-B07-B^T^ [4, 5] to 98.5 % with *Z. galactanivorans* Dsij^T^ [1] (Table S1). The complete genome of strain LLG6346-3.1^T^ was composed of 5,066,785 nucleotides and had a Chargaff’s coefficient of 38.28 %. Analysis of a phylogenomic tree based on 768 proteins from the core genome of sequenced *Zobellia* strains showed that LLG6346-3.1^T^ formed a clade with *Z. roscoffensis* Asnod1-F08^T^ (Fig. S2). The ANI and GGDC values for strain LLG6346-3.1^T^, when compared with other *Zobellia* species, were less than 90 % and less than 40 % respectively (88.0 % and 37.1 % with *Z. roscoffensis* Asnod1-F08^T^; Table S2). As the normally accepted thresholds of species delineation for ANI and GGDC are 95 % and 70%, respectively [29, 31, 37, 38], these values suggest that strain LLG6346-3.1^T^ represents a new species of the genus *Zobellia*.

**Fig. 1.**
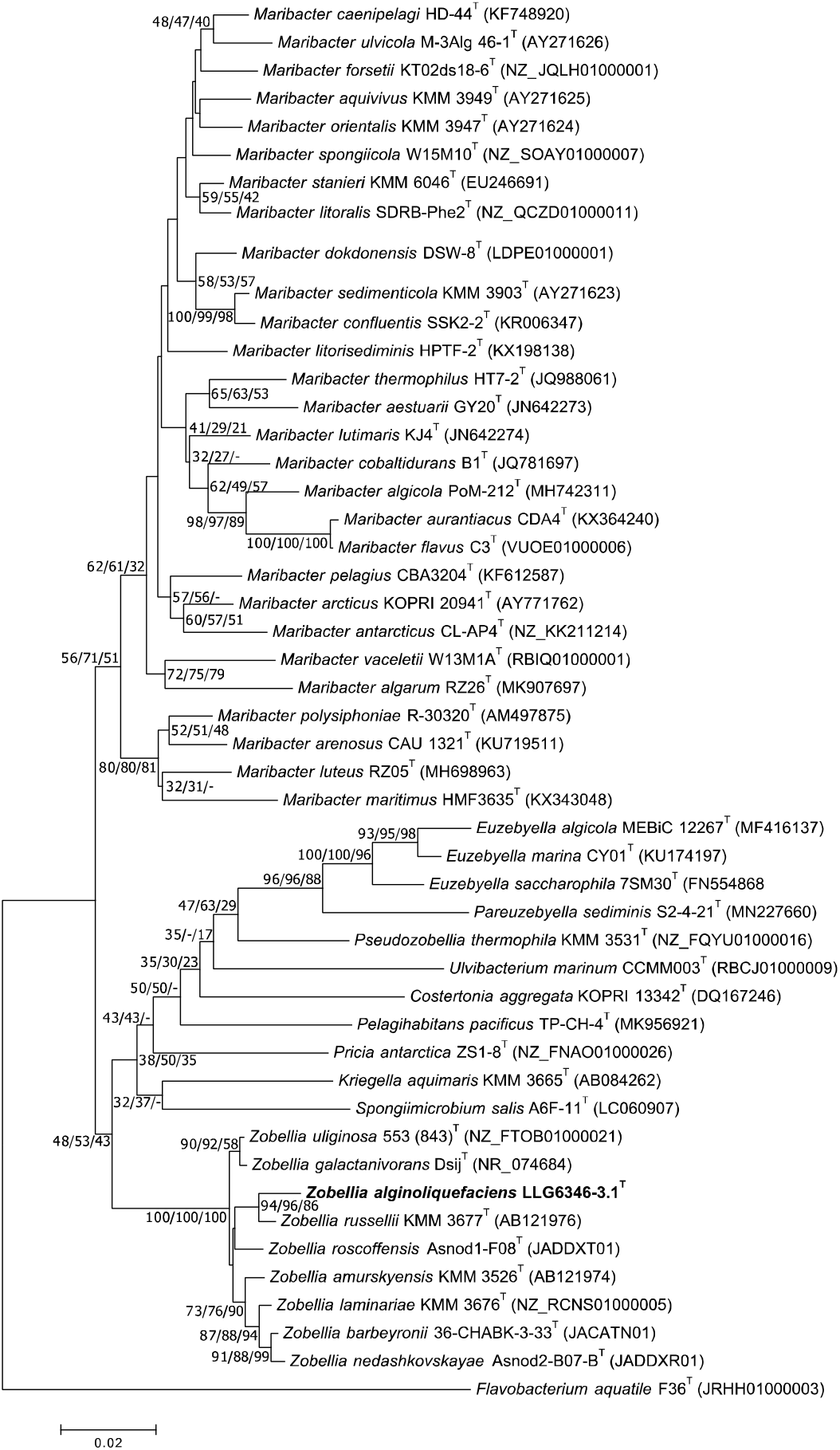
Neighbour-joining tree based on 16S rRNA gene sequences, showing the phylogenetic relationships between the strain LLG6346-3.1^T^ and related taxa from the family *Flavobacteriaceae*. Numbers at the nodes indicate bootstrap values (in percentage of 1000 replicates) from neighbour-joining, maximum-likelihood and maximum-parsimony analyses respectively, while dashes instead of numbers indicate that the node was not observed in the corresponding analysis. For nodes conserved in at least two trees, all bootstrap values are shown. Nodes without bootstrap value are not conserved in other trees and <70%. *Flavobacterium aquatile* F36^T^ was used as an outgroup. Bar, 0.02 changes per nucleotide position.

Under the microscope, cells of strain LLG6346-3.1^T^ appeared as rods approximately 0.5 μm in diameter and 2.0–4.0 μm long, attached to the glass of the slide or coverslip and showed gliding motility. Colonies grown on R2A agar at 20 °C showed a weak iridescence. The optimum growth temperature and NaCl concentration for strain LLG6346-3.1^T^ were higher than for *Z. russellii* KMM 3677^T^ and *Z. roscoffensis* Asnod1-F08^T^ (Table 1). This could reflect adaptation to the Mediterranean Sea environment from which strain LLG6346-3.1^T^ was isolated, where average seawater temperature and salinity are higher than the Pacific Ocean and English Channel from which *Z. russellii* KMM 3677^T^ and *Z. roscoffensis* Asnod1-F08^T^ were retrieved, respectively.

**Table 1.** Phenotypic characteristics of strain LLG6346-3.1^T^ and of two *Zobellia* species used as related type strains Strains: 1, LLG6346-3.1 (*Zobellia alginoliquefaciens* sp. nov.); 2, *Z. roscoffensis* Asnod1-F08^T^; 3, *Z. russellii* KMM 3677^T^. Cells of all strains share the following characteristics: Gram-negative, aerobic, heterotroph, chemorganotroph with respiratory metabolism, gliding motility, do not form endospores, do not accumulate poly-ß-hydroxybutyrate as an intracellular reserve product; require Na^+^ ion or seawater for growth and produce flexirubin-type pigments. All strains are positive for the utilization as a sole carbon source of D-glucose, D-galactose, D-fructose, D-mannose, D-xylose, salicine (weakly), sucrose, D-maltose, lactose, D-cellobiose, gentiobiose, trehalose, raffinose, D-mannitol, N-acetyl-glucosamine, 1-O-methyl-D-glucoside, 1-O-methyl-D-mannoside, lichenin, galactan (gum arabic), glucomannan, xylan and porphyrin; for the hydrolysis of DNA, aesculin, gelatin and Tween 20; for acid and alkaline phosphatase, esterase (C 4), esterase lipase (C 8), leucine arylamidase, valine arylamidase, cystine arylamidase, trypsine, naphtol AS-BI-phosphohydrolase, α-glucosidase, β-glucosidase, α-galactosidase, β-galactosidase (PNPG and API ZYM tests), N-acetyl-β-glucosaminidase, α-mannosidase, oxidase and catalase activities; for the acid production from D-glucose, D-galactose, D-fructose, D-mannose, D- and L-arabinose, D-xylose, D-lyxose, D-tagatose, D- and L-fucose, salicine (weakly), D-mannitol, 1-O-methyl-D-glucoside, 1-O-methyl-D-mannoside, D-maltose, lactose, melibiose, sucrose, D-cellobiose, D-turanose, trehalose, amygdaline, raffinose, and arbutine. All strains are negative for indole and H_2_S production; for utilization as a sole carbon source of D-arabinose, D-fucose, ribose, L-sorbose, L-xylose, D-lyxose, D-sorbitol, dulcitol, inositol, adonitol, erythritol, xylitol, D- and L-arabitol, arbutine, D-tagatose, gluconic acid, citric acid, capric acid, adipic acid, malic acid, phenylacetic acid, 2 ketogluconate, 5 ketogluconate, arabinan, fucoidin from *Ascophyllum nodosum*, fucoidin from *Laminaria hyperborea*, galactomannan, agar, κ-, ι- and λ-carrageenin, pectin, ulvin and xyloglucan; for the hydrolysis of Tween 40, κ- and ι-carrageenin; for urease, arginine dihydrolase, tryptophan deaminase, lysine decarboxylase, ornithine decarboxylase lipase (C 14), α-chemotrypsine, β-glucuronidase and α-fucosidase activities; for the acid production from L-xylose, glycerol, erythritol, adonitol, dulcitol, D-sorbitol, xylitol, D- and L-arabitol, N-acetyl-glucosamine, gluconic acid and 5 ketogluconate. +, Positive; -, negative; w, weakly positive; na, not available; MMM, Marine Minimum Medium; (liq.), liquefaction.

| Characteristic | 1 | 2 | 3 |
| --- | --- | --- | --- |
| Growth conditions: |  |  |  |
| Temperature range | 4-38 | 4-40* | 4-38<sup>#</sup> |
| Optimum temperature (°C) | 30-33 | 25-30* | 25-28<sup>#</sup> |
| pH range | 6.5-9 | 5.5-8.5* | na |
| Optimum pH | 8-8.5 | 7.5* | na |
| NaCl range (%) | 2-7 | 2-6* | 1-10<sup>#</sup> |
| Optimum NaCl (%) | 4-5 | 2* | 2-3<sup>#</sup> |
| Enzyme: |  |  |  |
| Nitrate reductase (nitrated ZoBell medium) | w | + | + |
| Utilisation of (API20NE): |  |  |  |
| L-Arabinose | + | - | + |
| Utilisation of (API50CH): |  |  |  |
| L-Rhamnose | - | + | + |
| Melezitose | + | - | + |
| L-Fucose | - | + | + |
| Glycerol | - | - | + |
| 1-O-methyl-D-Xyloside | - | + | + |
| D-Turanose | + | + | - |
| Melibiose | + | + | - |
| Amygdaline | - | + | - |
| Starch | + | - | + |
| Glycogene | + | - | - |
| Utilisation of (MMM) |  |  |  |
| L-Arabinose | W | - | + |
| L-Rhamnose | - | + | + |
| Arabinoxylane | - | - | + |
| Alginic acid | + | - | + |
| Laminarin | - | + | + |
| Acid production (API 50CH): |  |  |  |
| Ribose | - | - | + |
| L-Sorbose | - | + | + |
| L-Rhamnose | - | + | + |
| Melezitose | + | - | ± |
| 1-O-methyl-D-Xyloside | - | - | + |
| Gentiobiose | + | - | - |
| Inositol | - | - | + |
| Starch | + | - | + |
| Glycogene | + | - | - |
| 2 Ketogluconate | + | + | - |
| Hydrolysis of: |  |  |  |
| Starch (lugol assay) | + | - | + |
| Agar (lugol assay) | - | + | + |
| Alginic acid | + (liq.) | - | + |
| DNA G+C (mol%) | 38.28 | 37.61 | 39.03 |

\*Data from Barbeyron *et al.* [5]
#Data from Nedashkovskaya *et al.* [3]

Growth was observed with some polysaccharides and a few simple sugars allowing differentiation of strain LLG6346-3.1^T^ from *Z. russellii* KMM 3677^T^ and *Z. roscoffensis* Asnod1-F08^T^ (Table 1). The most obvious test to differentiate strain LLG6346-3.1^T^ from *Z. russellii* KMM 3677^T^ and *Z. roscoffensis* Asnod1-F08^T^ was hydrolysis of alginic acid. While strain LLG6346-3.1^T^ hydrolysed to liquefaction and used this marine polysaccharide as the sole carbon source, it was not hydrolysed and therefore not utilized by *Z. roscoffensis* Asnod1-F08^T^. For its part, *Z. russellii* KMM 3677^T^ hydrolysed, used but did not liquefy alginic acid, and only the formation of a crater in the alginic acid, without liquid, was visible (Table 1). Comparative genomics within the genus *Zobellia* suggests that the ability to hydrolyse alginic acid is mainly due to the presence of the alginate lyase-encoding genes *alyA1* and *alyA2* (zgal_1182 and zgal_2618 in genome of *Z. galactanivorans* Dsij^T^ respectively). The LLG6346-3.1^T^ strain, which liquefied alginic acid, possesses both genes. *Z. russellii* KMM 3677^T^ that hydrolysed alginic acid without liquefaction, possesses only the *alyA2* gene. By contrast *Z. roscoffensis* Asnod1-F08^T^, although possessing homologs of *alyA3, alyA4, alyA5* and *alyA6* genes from *Z. galactanivorans* Dsij^T^, does not possess either the *alyA1* or the *alyA2* genes. These observations suggest that the liquefaction phenotype is linked to the presence of the *alyA1* gene encoding a secreted endo-guluronate lyase [39]. Among valid species of *Zobellia*, only *Z. galactanivorans* Dsij^T^, *Z. uliginosa* 553(843)^T^ [1] and *Z. nedashkovskayae* Asnod2-B07-B^T^ [5] possess the *alyA1* gene and liquefied and utilized alginic acid. However, it is easy to differentiate these species. Unlike *Z. galactanivorans* Dsij^T^ which is able to hydrolyse all red algal polysaccharides (agars and carrageenins) and *Z. nedashkovskayae* Asnod2-B07-B^T^ which is able to utilize laminarin and fucoidin from *Ascophyllum nodosum*, strain LLG6346-3.1^T^ did not hydrolyse and use neither agars nor carrageenins (which is consistent with the absence of carrageenase-encoding genes in its genome) nor laminarin nor fucoidin (Table 1). Strains LLG6346-3.1^T^ and *Z. russellii* KMM 3677^T^ were able to use starch as sole carbon and energy sources and showed a hydrolysis area on soluble starch ZoBell agar plates (Table 1). This suggests that both strains possess a secreted alpha-amylase, consistent with the presence of the amylase-encoding gene *susA* in their genome. As reported previously [5], *Z. roscoffensis* Asnod1-F08^T^ lacks a *susA* homolog, likely explaining its absence of hydrolysis and use of starch. Finally, the nitrate reductase activity is another discriminant criteria to differentiate the strain LLG6346-3.1^T^ from *Z. roscoffensis* Asnod1-F08^T^ and *Z. russellii* KMM 3677^T^. While the latter two strains showed vigorous nitrate reductase activity after growth in nitrated ZoBell broth, strain LLG6346-3.1^T^ showed very weak activity under the same conditions.

The other physiological features of strain LLG6346-3.1^T^ compared with *Z. roscoffensis* Asnod1-F08^T^ and *Z. russellii* KMM 3677^T^ are listed in Table 1. The three strains were resistant to kanamycin, gentamycin, neomycin, vancomycin, ampicillin, penicillin, carbenicillin, oxacillin, erythromycin, nalidixic acid, trimethoprim/sulfamethoxazole, bacitracin, colistin, polymixin B and chloramphenicol. For streptomycin, whereas *Z. roscoffensis* Asnod1-F08^T^ is sensible, the other two strains are resistant. In the case of lincomycin, whereas the strain LLG6346-3.1^T^ is sensitive, the other two strains are resistant. Finally, the strain LLG6346-3.1^T^, *Z. roscoffensis* Asnod1-F08^T^ and *Z. russellii* KMM 3677^T^ were sensitive to rifampicin. In conclusion, phenotypic characterizations and phylogenetic analysis using 16S rRNA gene sequences together with whole-genome pairwise comparisons show that strain LLG6346-3.1^T^ represents a novel species in the genus *Zobellia*, for which the name *Zobellia alginoliquefaciens* sp. nov. is proposed.

## DESCRIPTION OF *ZOBELLIA ALGINOLIQUEFACIENS* SP. NOV

*Zobellia alginoliquefaciens* (al.gi.no.li.que.fa’ci.ens. N.L. pres. part. *liquefaciens*, liquefying; N.L. part. adj. *alginoliquefaciens*, alginic acid digesting).

Cells are Gram-stain-negative, aerobic, chemoorganotrophic, heterotrophic and rod-shaped, approximately 0.5 μm in diameter and 2.0–4.0 μm long; a few cells greater than 4 μm long may occur. Flagella are absent. Prosthecae and buds are not produced. Colonies on ZoBell agar are orange-coloured, convex, circular and mucoid in consistency and 2.0–3.0 mm in diameter after incubation for 3 days at 20 °C. Growth in ZoBell 2216E broth occurs from 4 to 38 °C (optimum, 30-33°C), at pH 6.5–9.0 (optimum, pH 8-8.5) and in the presence of 2–7% NaCl (optimum, 4-5%). Positive for gliding motility and flexirubin-type pigment production. Nitrate is very weakly reduced. β-Galactosidase-, oxidase- and catalase-positive. Alginic acid is hydrolysed to total liquefaction. DNA, gelatin, starch, aesculin, Tweens 20 and Tweens 60 are hydrolysed, but Tween 40, agar, κ-carrageenin and ι-carrageenin are not. D-glucose, D-galactose, D-fructose, L-arabinose, D-mannose, D-xylose, salicin (weakly), sucrose, lactose, D-maltose, melibiose, D-cellobiose, gentiobiose, D-turanose, trehalose, D-mannitol, melezitose, raffinose, N-acetyl-glucosamine, starch, glycogene, inulin (weakly), porphyrin, alginic acid, xylan, galactan (gum arabic), glucomannan, lichenin, 1-O-methyl-D-glucoside and 1-O-methyl-D-mannoside are utilized as carbon and energy sources, but D-arabinose, ribose, L-fucose, D-fucose, D-lyxose, L-rhamnose, L-sorbose, D-tagatose, L-xylose, arbutine, amygdaline, adipic acid, capric acid, malic acid, citric acid, gluconic acid, phenylacetic acid, 2 ketogluconate, 5 ketogluconate, 1-O-methyl-D-xyloside, adonitol, D-arabitol, L-arabitol, dulcitol, erythritol, glycerol, inositol, D-sorbitol, xylitol, arabinan, arabinoxylane, pectin (apple), galactomannan, xyloglucan, agar, ι-carrageenin, κ-carrageenin, λ-carrageenin, laminarin, ulvin, fucoïdin (*Ascophyllum nodosum*) and fucoïdin (*Laminaria hyperborea*) are not. Acid is produced from D-glucose, D-galactose, D-tagatose, D-fructose, D-arabinose, L-arabinose, D-mannose, D-fucose, L-fucose, D-lyxose, D-xylose, salicin (weakly), arbutine, sucrose, lactose, D-maltose, melibiose, D-cellobiose, gentiobiose, D-turanose, trehalose, amygdaline, melezitose, raffinose, starch, glycogene, inulin (weakly), D-mannitol, 1-O-methyl-D-glucoside, 1-O-methyl-D-mannoside, and 2 ketogluconate, but not from ribose, L-xylose, L-sorbose, L-rhamnose, glycerol, erythritol, inositol, D-sorbitol, dulcitol, xylitol, D-arabitol, L-arabitol, adonitol, N-acetyl-glucosamine, 5 ketogluconate and 1-O-methyl-D-xyloside. Negative for indole and H_2_S production and for arginine dihydrolase, tryptophan deaminase, urease, lysine decarboxylase and ornithine decarboxylase activities. In the API ZYM system, activities from acid phosphatase, alkaline phosphatase, esterase (C4), esterase lipase (C8), leucine arylamidase, valine arylamidase, cystine arylamidase, trypsin, naphthoL-AS-BI-phosphohydrolase α-galactosidase, ß-galactosidase, α-glucosidase, ß-glucosidase, N-acetyl-ß-Glucosaminidase and α-Mannosidase are present, but activities of lipase (C14), α-chymotrypsin, ß-glucuronidase and α-fucosidase are absent. The DNA G+C content is 38.28%.

The type strain, LLG6346-3.1^T^ (RCC 7657^T^ = LLG 32918^T^), was isolated from the surface of the brown alga *Ericaria zosteroides* (C.Agardh) Molinari & Guiry 2020.

The Genbank/EMBL/DDBJ accession number for the 16S rRNA gene sequence of strain LLG6346-3.1^T^ is OQ511313. The Genbank/EMBL/DDBJ accession number for the genome sequence of strain LLG6346-3.1^T^ is SAMN33610405.

## Supporting information

Strain deposit certificate 1

Strain deposit certificate 2

## Abbreviations

ANI: average nucleotide identity
dDDH: digital DNA-DNA hybridization
GGDC: Genome-Genome Distance Calculator
ML: maximum-likelihood
MP: maximumparsimony
NJ: neighbour-joining

## Acknowledgements

The LABGeM (CEA/Genoscope and CNRS UMR8030), the France Génomique and French Bioinformatics Institute national infrastructures (funded as part of Investissement d’Avenir program managed by Agence Nationale pour la Recherche, contracts ANR-10-INBS-09 and ANR-11-INBS-0013) are acknowledged for support within the MicroScope annotation platform. We are grateful to the Institut Français de Bioinformatique (ANR-11-INBS-0013) and the Roscoff Bioinformatics platform ABiMS (http://abims.sb-roscoff.fr) for providing help for computing and storage resources. This work has benefited from the facilities of the Genomer platform, which is part of the Biogenouest core facility network. We thank Gwenn Tanguy and Erwan Legeay for sequencing. This work has benefited from the CORSICABENTHOS expeditions (PI: Line Le Gall), the marine component of the “*Our Planet Reviewed*” programme. The Corsica programme is run by Muséum National dHistoire Naturelle in partnership with Université de Corse Pasquale Paoli and Office de l’Environnement de la Corse (OEC), with the support of Office Français de la Biodiversité (OFB) and Collectivité Territoriale de Corse (CTC). The CORSICABENTHOS 1 expedition took place in May 2019 in collaboration with Parc Naturel Marin du Cap Corse et de l’Agriate ; The organizers are grateful to Madeleine Cancemi, Jean-François Cubells, Jean-Michel Culioli and Jean-Michel Palazzi for their support.

---

**Supplementary Table S1:** Assembly statistics and properties of strain LLG6346-3.1^T^ complete genome

|  | **LLG6346-3.1<sup>T</sup>** |
| --- | --- |
| **Illumina sequencing** |  |
| Paired sequencing reads | 4,268,034 |
| Total sequencing data | 1,284,678,234 nt |
| **Nanopore sequencing** |  |
| Paired sequencing reads | 100,270 |
| Total sequencing data | 2,299,667,950 nt |
| average read length | 22,935 nt |
| **Assembled complete genome** |  |
| genome accession |  |
| length | 5,066,785 nt |
| estimated coverage | 707 X |
| G+C content | 38.28 |
| number of CDS | 4,143 |

**Supplementary Table S2.**
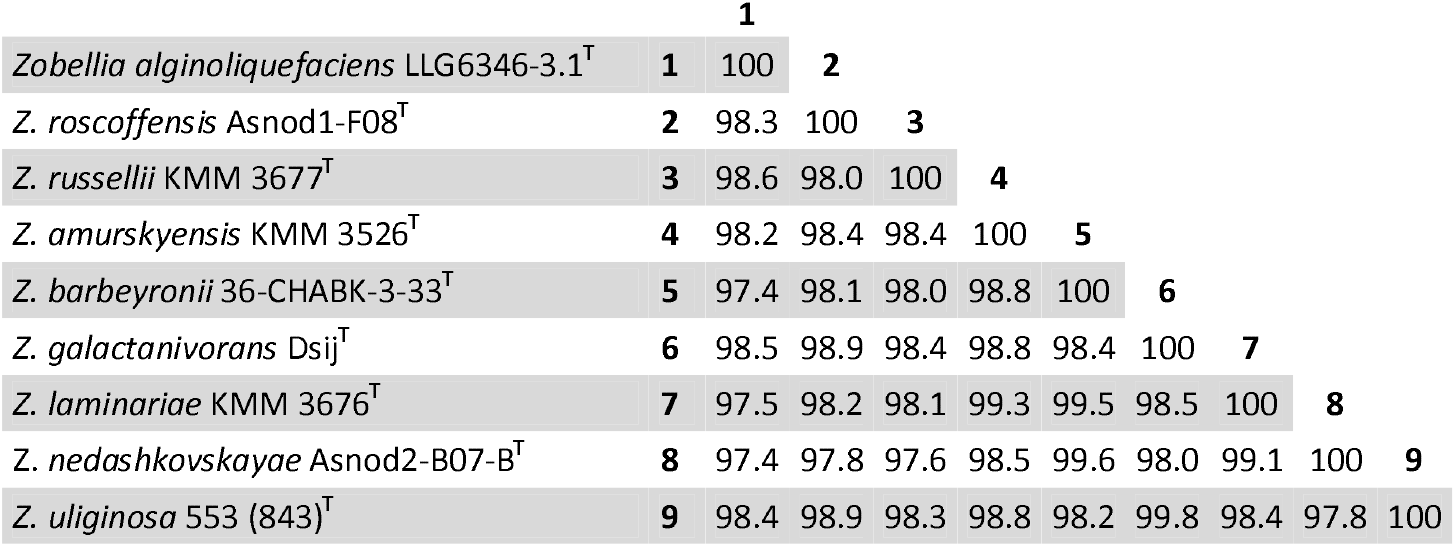
Pairwise nucleotide comparisons (in percentage of similarity sequence) between *Zobellia* species 16S rRNA gene sequences.

**Supplementary Table S3.**
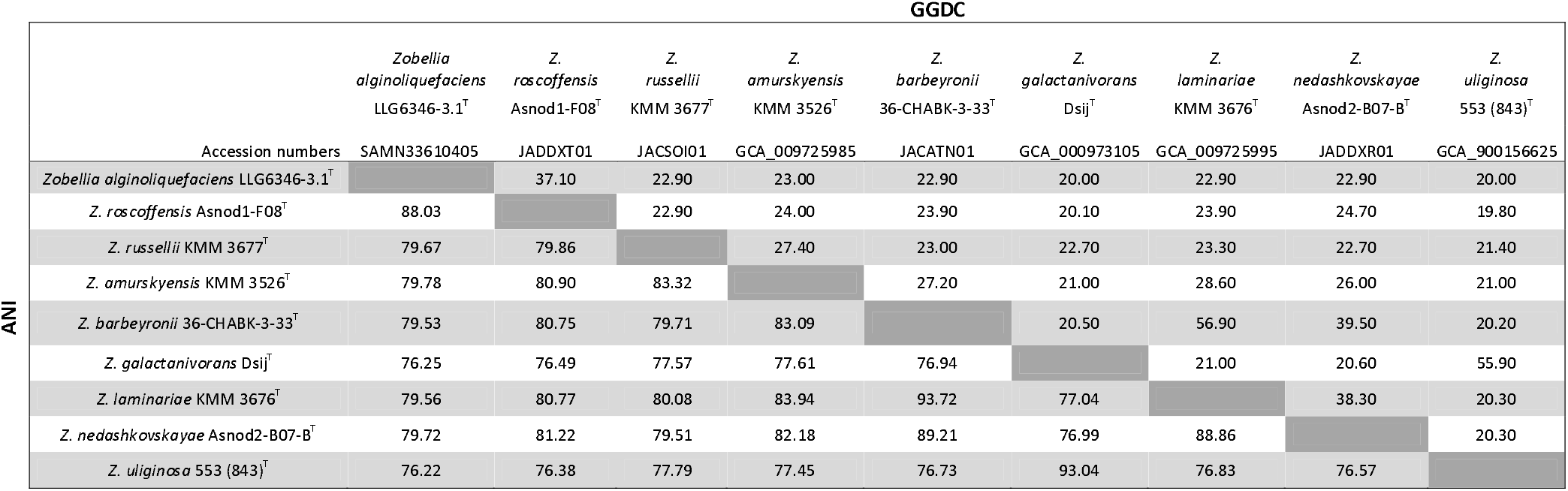
Whole genome relatedness analysis (in percentage) of the strain LLG6346-3.1^T^ and *Zobellia* species. Average nucleotide identities (ANI) are shown below the diagonal and genome-to-genome distance calculations (GGDC) are shown above the diagonal. GGDC values were calculated online (http://ggdc.dsmz.de/distcalc2.php) and the results from formula 2 were used. ANI values were calculated with an online program (http://jspecies.ribohost.com/jspeciesws/#analyse). The GenBank accession numbers for the whole genome sequences are given in the top row.

**Supplementary Fig. S1.**
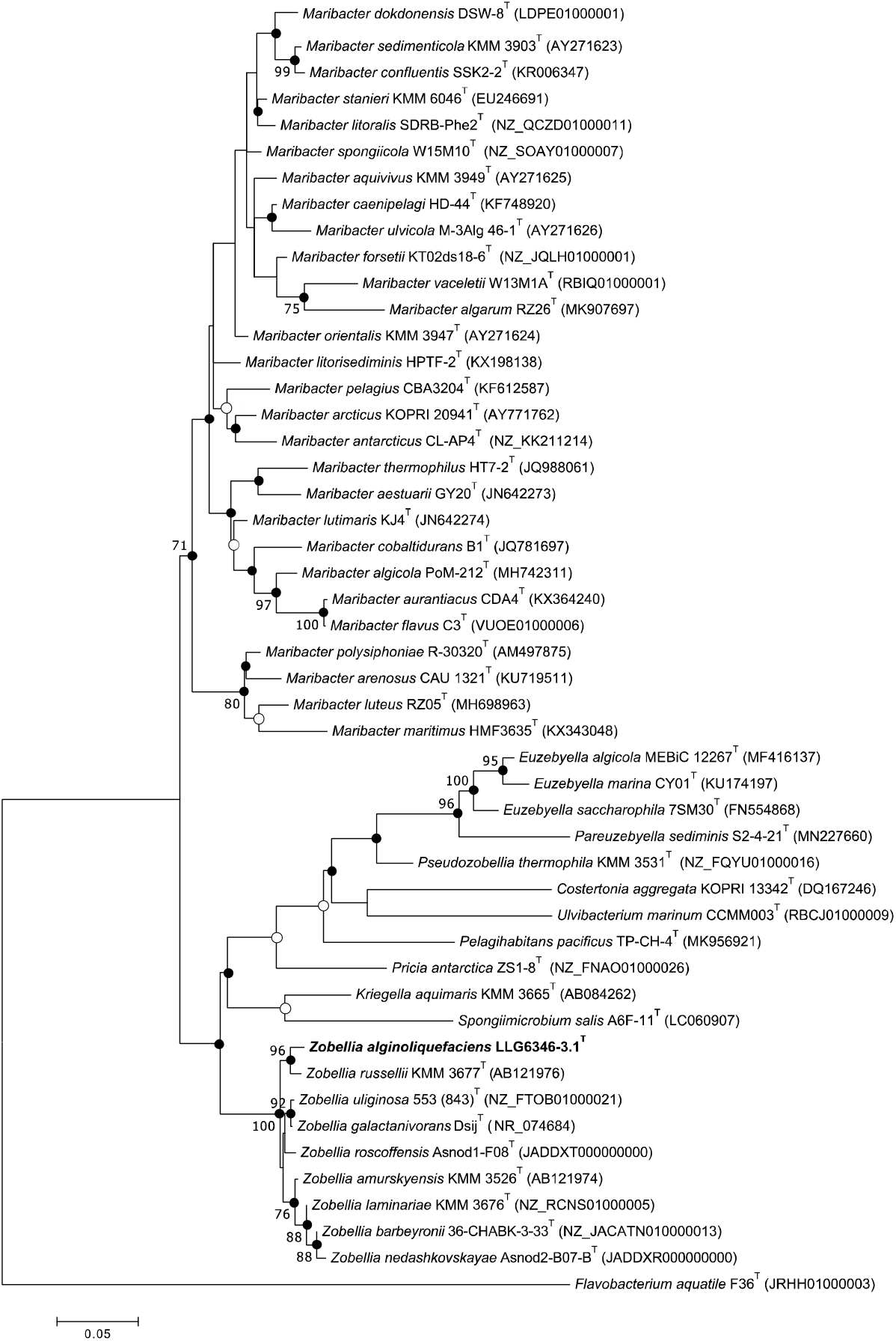
Maximum-likelihood (ML) tree based on 16S rRNA gene sequences, showing the phylogenetic relationships between the strain LLG6346-3.1^T^ and related taxa from the family *Flavobacteriaceae*. Numbers at nodes are bootstrap values shown as percentages of 1000 replicates; only values >70 % are shown. Filled circles indicate that the corresponding nodes were also recovered in the trees generated with the neighbour-joining (NJ) and maximumparsimony algorithms, while open circles indicate that the nodes were only recovered in the tree generated with the NJ and ML algorithms. *Flavobacterium aquatile* F36^T^ was used as an outgroup. Bar, 0.05 changes per nucleotide position.

**Supplementary Fig. S2.**
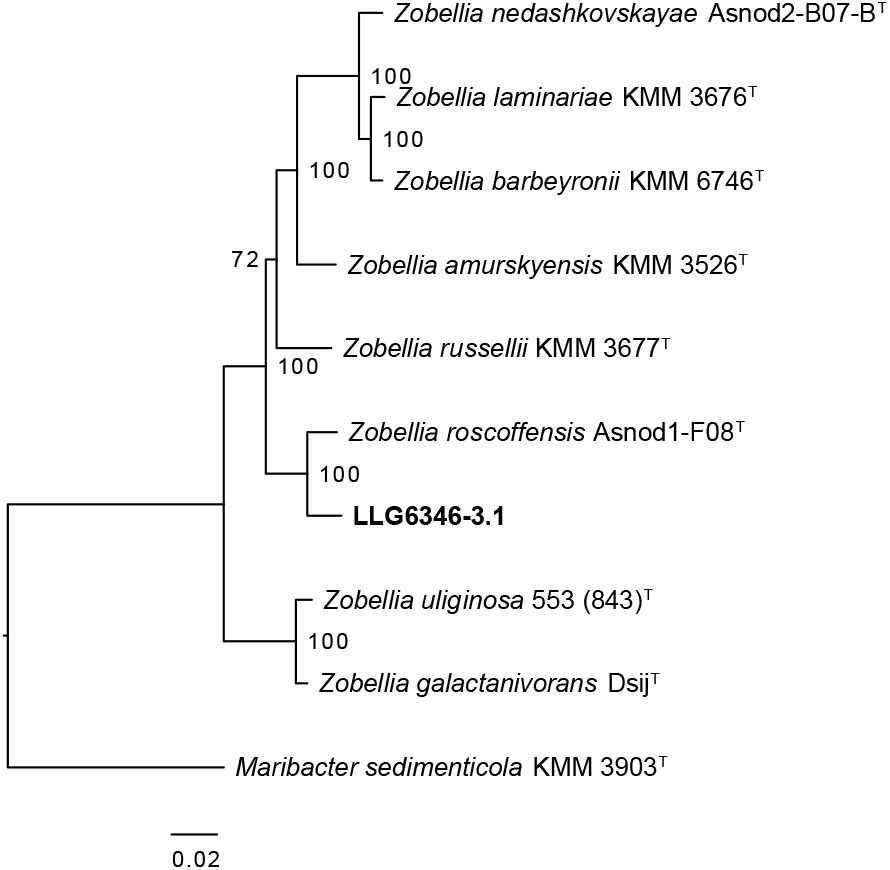
Core proteome phylogenetic analysis of available genomes from *Zobellia* type strains, the newly isolated LLG6346-3.1^T^ strain and *Maribacter sedimenticola* KMM 3903^T^ used as an outgroup. Bar, 0.02 substitutions per amino acid position.

## References

1. Barbeyron T, L’Haridon S, Corre E, Kloareg B, Potin P. *Zobellia galactanovorans* gen. nov., sp. nov., a marine species of *Flavobacteriaceae* isolated from a red alga, and classification of [*Cytophaga*] *uliginosa* (ZoBell and Upham 1944) Reichenbach 1989 as *Zobellia uliginosa* gen. nov., comb. nov. Int J Syst Evol Microbiol 2001;51:985–97.

2. Potin P, Sanseau A, Le Gall Y, Rochas C, Kloareg B. Purification and characterization of a new k-carrageenase from a marine *Cytophaga*-like bacterium. Eur J Biochem 1991;201:241–247.

3. Nedashkovskaya OI, Suzuki M, Vancanneyt M, Cleenwerck I, Lysenko AM, et al. *Zobellia amurskyensis* sp. nov., *Zobellia laminariae* sp. nov. and *Zobellia russellii* sp. nov., novel marine bacteria of the family *Flavobacteriaceae*. Int J Syst Evol Microbiol 2004;54:1643–8.

4. Nedashkovskaya O, Otstavnykh N, Zhukova N, Guzev K, Chausova V, et al. *Zobellia barbeyronii* sp. nov., a New Member of the Family *Flavobacteriaceae,*Isolated from Seaweed, and Emended Description of the Species *Z. amurskyensis*, *Z. laminariae*, *Z. russellii* and *Z. uliginosa*. Diversity 2021;13:520.

5. Barbeyron T, Thiébaud M, Le Duff N, Martin M, Corre E, et al. *Zobellia roscoffensis* sp. nov. and *Zobellia nedashkovskayae* sp. nov., two flavobacteria from the epiphytic microbiota of the brown alga *Ascophyllum nodosum*, and emended description of the genus *Zobellia*. Int J Syst Evol Microbiol;71. Epub ahead of print 2021. DOI: 10.1099/ijsem.0.004913.

6. Miranda LN, Hutchison K, Grossman AR, Brawley SH. Diversity and abundance of the bacterial community of the red macroalga *Porphyra umbilicalis:* Did bacterial farmers produce macroalgae? PLoS One 2013;8:e58269.

7. Dogs M, Wemheuer B, Wolter L, Bergen N, Daniel R, et al. *Rhodobacteraceae* on the marine brown alga *Fucus spiralis* are abundant and show physiological adaptation to an epiphytic lifestyle. Syst Appl Microbiol 2017;40:370–382.

8. Brunet M, Le Duff N, Fuchs BM, Amann R, Barbeyron T, et al. Specific detection and quantification of the marine flavobacterial genus *Zobellia* on macroalgae using novel qPCR and CARD-FISH assays. Syst Appl Microbiol 2021;44:126269.

9. Zobell C. Studies on marine bacteria I The cultural requirements of heterotrophic aerobes. J Mar Res 1941;4:42–75.

10. Brewster JD. A simple micro-growth assay for enumerating bacteria. J Microbiol Methods 2003;53:77–86.

11. Powers EM. Efficacy of the Ryu nonstaining KOH technique for rapidly determining gram reactions of food-borne and waterborne bacteria and yeasts. Appl Environ Microbiol 1995;61:3756–3758.

12. Smibert R, Krieg N. General characterization. In: Gerhardt P, Murray R, Costilow R, Nester E, Wood W, et al. (editors). Manual of Methods for General Bacteriology. Washington, DC., USA: American Society for Microbiology; 1981. pp. 409–443.

13. Draget KI, Ostgaard K, Smidsrød O. Alginate-based solid media for plant tissue culture. Appl Microbiol Biotechnol 1989;31:79–83.

14. Thomas F, Barbeyron T, Michel G. Evaluation of reference genes for real-time quantitative PCR in the marine flavobacterium *Zobellia galactanivorans*. J Microbiol Methods;84. Epub ahead of print 2011. DOI: 10.1016/j.mimet.2010.10.016.

15. Wick RR, Judd LM, Gorrie CL, Holt KE. Unicycler: Resolving bacterial genome assemblies from short and long sequencing reads. PLoS Comput Biol 2017;13:1–22.

16. Hicks R, Amann R, Stahl D. Dual staining of natural bacterioplankton with 4’,6-diamidino-2-phenylindole and fluorescent oligonucleotide probes targeting kingdom-level 16S rRNA sequences. Appl Environ Microbiol 1992;58:2158–2163.

17. Kane M, Poulsen L, Stahl D. Monitoring the enrichment and isolation of sulfate-reducing bacteria by using oligonucleotide hybridization probes designed from environmentally derived 16S rRNA sequences. Appl Environ Microbiol 1993;59:682–686.

18. Katoh K. MAFFT: a novel method for rapid multiple sequence alignment based on fast Fourier transform. Nucleic Acids Res 2002;30:3059–3066.

19. Saitou N, Nei M. The neighbor-joining method: a new method for reconstructing phylogenetic trees. Mol Biol Evol 1987;4:406–425.

20. Fitch WM. Toward defining the course of evolution: Minimum change for a specific tree topology. Syst Biol 1971;20:406–416.

21. Felsenstein J. Evolutionary trees from DNA sequences: A maximum likelihood approach. J Mol Evol 1981;17:368–376.

22. Tamura K, Stecher G, Peterson D, Filipski A, Kumar S. MEGA6: Molecular evolutionary genetics analysis version 6.0. Mol Biol Evol 2013;30:2725–2729.

23. Kimura M. A simple method for estimating evolutionary rates of base substitutions through comparative studies of nucleotide sequences. J Mol Evol 1980;16:111–120.

24. Nei M, Kumar S. Molecular Evolution and Phylogenetics. New York: Oxford University Press.; 2000.

25. Felsenstein J. Confidence limits on phylogenies: an approach using the bootstrap. Evolution (N Y) 1985;39:783–791.

26. Avram O, Rapoport D, Portugez S, Pupko T. M1CR0B1AL1Z3R - a user-friendly web server for the analysis of large-scale microbial genomics data. Nucleic Acids Res 2019;47:W88–W92.

27. Stamatakis A. RAxML version 8: A tool for phylogenetic analysis and post-analysis of large phylogenies. Bioinformatics 2014;30:1312–1313.

28. Yoon SH, Ha SM, Kwon S, Lim J, Kim Y, et al. Introducing EzBioCloud: A taxonomically united database of 16S rRNA gene sequences and whole-genome assemblies. Int J Syst Evol Microbiol 2017;67:1613–1617.

29. Goris J, Konstantinidis KT, Klappenbach JA, Coenye T, Vandamme P, et al. DNA-DNA hybridization values and their relationship to whole-genome sequence similarities. Int J Syst Evol Microbiol 2007;57:81–91.

30. Figueras MJ, Beaz-Hidalgo R, Hossain MJ, Liles MR. Taxonomic affiliation of new genomes should be verified using average nucleotide identity and multilocus phylogenetic analysis. Genome Announc 2014;2:2–3.

31. Chun J, Oren A, Ventosa A, Christensen H, Arahal DR, et al. Proposed minimal standards for the use of genome data for the taxonomy of prokaryotes. Int J Syst Evol Microbiol 2018;68:461–466.

32. Auch AF, von Jan M, Klenk HP, Göker M. Digital DNA-DNA hybridization for microbial species delineation by means of genome-to-genome sequence comparison. Stand Genomic Sci 2010;2:117–134.

33. Meier-Kolthoff JP, Auch AF, Klenk HP, Göker M. Genome sequence-based species delimitation with confidence intervals and improved distance functions. BMC Bioinformatics;14. Epub ahead of print 2013. DOI: 10.1186/1471-2105-14-60.

34. Meier-Kolthoff JP, Göker M, Spröer C, Klenk HP. When should a DDH experiment be mandatory in microbial taxonomy? Arch Microbiol 2013;195:413–418.

35. Vallenet D, Belda E, Calteau A, Cruveiller S, Engelen S, et al. MicroScope - An integrated microbial resource for the curation and comparative analysis of genomic and metabolic data. Nucleic Acids Res 2012;41:1–12.

36. Lombard V, Golaconda Ramulu H, Drula E, Coutinho PM, Henrissat B. The carbohydrate-active enzymes database (CAZy) in 2013. Nucleic Acids Res 2014;42:490–495.

37. Richter M, Rosselló-Móra R. Shifting the genomic gold standard for the prokaryotic species definition. Proc Natl Acad Sci U S A 2009;106:19126–19131.

38. Auch AF, Klenk HP, Göker M. Standard operating procedure for calculating genome-to-genome distances based on high-scoring segment pairs. Stand Genomic Sci 2010;2:142–148.

39. Thomas F, Lundqvist LCE, Jam M, Jeudy A, Barbeyron T, et al. Comparative characterization of two marine alginate lyases from *Zobellia galactanivorans* reveals distinct modes of action and exquisite adaptation to their natural substrate. J Biol Chem;288:23021–23037. DOI: 10.1074/jbc.M113.467217.

