## Supplementary material for "*Zobellia alginoliquefaciens* sp. nov., a new flavobacteria isolated from the epibiota of the brown alga *Ericaria zosteroides* (C.Agardh) Molinari & Guiry 2020": Strain deposit certificate 1

GREAT AT SMALL THINGS

BCCM DCG

BCCM IHEM

BCCM ITM

BCCM LMBP

BCCM MUCL

BCCM ULC

BCCM CC

### CERTIFICATE OF DEPOSIT

**This is to certify that the following microorganism has been deposited into the public BCCM/LMG Bacteria Collection and will be available for research purposes without restrictions:**

**LMG number:** **LMG 32918**

**Speciesname:** **Zobellia sp. nov.**

**Depositor:** **Thomas François, Station Biologique de Roscoff, UMR8227**

**Depositor no:** **LLG6346-3.1**

**The strain has been checked for viability and is preserved using one of the standard methods at BCCM/LMG. Authenticity was confirmed by the LMG quality control check as communicated with the depositor.**

**BCCM/LMG is not responsible for eventual discrepancies in characterization of the deposited material and its original description.**

07 February 2023

Vereecke Claudine

ir. Claudine Vereecke
Curator BCCM/LMG
