## Supplementary material for "*Zobellia alginoliquefaciens* sp. nov., a new flavobacteria isolated from the epibiota of the brown alga *Ericaria zosteroides* (C.Agardh) Molinari & Guiry 2020": Strain deposit certificate 2

Roscoff Culture Collection
Sorbonne University / CNRS
FR2424 Station Biologiquie de Roscoff
Place Georges Teissier
29682 Roscoff cedex
France

### Roscoff Culture Collection
### Strain Deposit Certificate

Depositor's family name: THOMAS
Depositor’s first name: François
Depositor’s position: Researcher
Depositor's institute name: CNRS Station Biologique de Roscoff
Depositor’s institute full address: Place Georges Teissier, 29682 Roscoff
Depositor’s

Date: 08/12/2022

**The RCC hereby acknowledges the deposit of 1 culture strain of Zobelia sp. (RCC 7657) in the Roscoff Culture Collection, as follows:**

- Zobellia sp. strain LLG6346-3.1 RCC7657

**According to RCC policy, these culture strain will henceforth be available to the scientific community without restriction of use via the on-line catalogue of the collection (www.roscoff-culture-collection.org).**

GOURVIL

Priscillia GOURVIL (RCC Curator)
